# miR-216b-5p is Down-Regulated in Human Breast Cancer and Inhibits Proliferation and Progression by Targeting HDAC8 Oncogene

**DOI:** 10.1101/499905

**Authors:** Mohammad-Nazir Menbari, Karim Rahimi, Abbas Ahmadi, Samira Mohammadi-Yeganeh, Anvar Elyasi, Nikoo Darvishi, Vahedeh Hosseini, Mohammad Abdi

## Abstract

**Background:** Previous studies showed the role of histone deacetylases (HDACs) on the progression of some malignancies. Recently, there is more attention to therapeutic applications of epigenetic factors such as microRNAs (miRs). To the best of our knowledge, there are no other results regarding the contribution of miR-216b-5p and its potential target, HDAC8, in progression of cancer.

**Aim:** In the present study, we investigated the role of miR-216b-5p on HDAC8 and there following impacts on breast cancer (BC) progression.

**Methods:** Human BC specimens and noncancerous tissues were acquired from Iran Tumor Bank (I.T.B). The MDA-MB-231, MCF-7 and MCF-10A BC cell lines were also prepared. The tissue and cell line expression levels of miR-216b-5p and HDAC8 were determined by quantitative real-time PCR (qPCR). Protein levels of HDAC8 were also measured by Western blotting assay. The cell cycle, cell proliferation and colony formation assay were determined and the role of HDAC8 was investigated using a knockout vector. Targeting the 3′ untranslated region (3′UTR) of HDAC8 by miR-216b-5p were confirmed using a luciferase reporter assay.

**Results:** Our results show the significant decline in miR-216b-5p and highly increase in HDAC8 levels in human breast cancer tissues and cell lines. Overexpression of miR-216b-5p in BC cell lines inhibited cellular proliferation and progression and inhibition of miR-216b-5p reverse these effects. We also confirmed that HDAC8 is directly down-regulated by miR-216b-5p. Knockout of HDAC8 had also the effects as miR-216b-5p overexpression. Furthermore, we found that lower levels of miR-216b-5p are negatively correlated with lymph node metastasis and advanced tumor size.

**Conclusion:** Taken together, our data shown that targeted overexpression of miR-216b-5p can suppress the growth of BC cells down-regulation of HDAC8. Therefore, inhibition of HDAC8 by miR-216b-5p will be helpful developing newer therapies for the effective treatment of BC.

## Introduction

Breast cancer (BC) is the most common malignancy in women which accounts for 26% of all cancers by 182,000 new cases annually (Siegel et al., 2018). BC is a complicated and complex disease characterized by the growth of malignant cells in the mammary glands with a great number of aberrations at the pathologic, genomics, epigenetics, and molecular level which eventually results in the dysregulation of gene expression and signaling pathways (Gourd, 2018).

Aberrant expression of histone deacetylases (HDACs) have been related to tumor pathogenesis, progression, and prognosis (Zhang et al., 2017). Therefore, these enzymes are among the most potential biomarkers and therapeutic targets for cancer detection and treatment (Zhang et al., 2017). Recently, there has been more attention to HDAC inhibitors as potential therapeutic agents for treatment of some malignancies (Chakrabarti et al., 2015; Zhang et al., 2017). Among them, HDAC8 is the most recently identified class I HDACs which has been linked to a number of diseases particularly to hematological malignancies (Huang and Geng, 2017). Recent studies have shown that HDAC8 is a potential oncogene and its high expression is directly related to several malignancies (Chakrabarti et al., 2015). Besides, siRNA mediated down regulation of HDAC8 has shown anti-tumor activity of HDAC8 and expression inhibition of HDAC8 increases the ratio of apoptotic cells in T cell lymphomas (Moser et al., 2014). Additionally, it seems that HDAC8 involved in the pathogenesis of neuroblastoma (Oehme et al., 2009).

MicroRNAs (miRs) are natural RNA molecules that play crucial roles in cellular processes and regulate gene expression post-transcriptionally. MiRs are a class of small non-coding RNAs with 19-25 bp length cleaved from a 70-100 bp hairpin pre-miRNA precursors (Mohr and Mott, 2015). Single-stranded mature miR binds to the 3′ untranslated region (3′UTR) of the target mRNA which results in either the block of translation or degradation of targeted mRNA molecule (Mohr and Mott, 2015). Previous studies identified numerous miRs that their aberrant expression is associated with various human malignancies (Acunzo et al., 2015). Among them, miR-216b-5p has been proposed as a tumor suppressor in nasopharyngeal carcinoma which acts through the suppression of protein kinase C alpha (PKCα) or the Kirsten rat sarcoma viral oncogene homolog (KRAS) (Deng et al., 2011; Liu et al., 2015); however, the roles of miR-216b in the progression and development of breast cancer remain unclear.

The concurrent association between miR-216 and HDAC8 with progression of breast cancer is not understood yet. Besides, to the best of our knowledge, there is no other study with regard to the evaluation of the role of miR-216 and HDAC8 in the carcinogenesis of breast cancer. Hence, in the present study we aimed to study the miR-216b-5p and HDAC8 expressions in patients with BC and healthy subjects. Besides, the present study attempts to assess the potential role of miR-216 (as a potential inhibitor of the HDAC8) on tumor growth in human breast adenocarcinoma.

## Material and methods

### Patients and tissue collection

During January 2016 to January 2018, a total of 32 tissue specimens were obtained from Iran Tumor Bank (I.T.B). Control samples were obtained from adjacent normal tissue and both malignant and normal tissues were confirmed pathologically. Written informed consent was obtained from all patients, and the study has been approved by the ethics committee of Kurdistan University of Medical Sciences. Patients with a history of other organ cancers were excluded from the study. The Scarf–Bloom–Richardson criteria and the TNM staging system for breast cancer were used for tumor grading and evaluating the clinical staging of patients, respectively (Edge et al., 2010; Elston, 2005). We obtained information on morphologic characteristics, grade, and stage of the tumor from the medical records (Table 1).

**Table 1:** Clinical characteristics of studied subjects.

| **Table 1: Clinical characteristics of studied subjects** |  |  |  |  |  |  |  |  |  |  |
| --- | --- | --- | --- | --- | --- | --- | --- | --- | --- | --- |
| Sample | site | Type of breast cancer | Stage of disease | Expression of hormone receptors |  |  | Necrosis | Invasion to other tissues |  |  |
|  |  |  |  | ER | PR | Her2/neu |  | Lymphatic invasion | Vascular invasion | perineural invasion |
| 1 | Right breast | infiltrating ductal carcinoma | III | Neg | Neg | Neg | Yes | Yes | Yes | No |
| 2 | Right breast | invasive ductal carcinoma | III | Neg | Neg | Neg | Yes | Yes | No | No |
| 3 | Left breast |  | III | Neg | Neg | Neg | Yes | Yes | Yes |  |
| 4 | Right breast | invasive lobular carcinoma | I | Neg | Neg | Neg | No | Yes | No | Yes |
| 5 | Left breast |  | III | Neg | Neg | Neg | Yes | Yes | Yes |  |
| 6 | Right breast | invasive ductal carcinoma | II | Neg | Neg | Neg | No | No | No | No |
| 7 | Left breast | invasive ductal carcinoma | II | Neg | Neg | Neg | Yes | Yes | Yes | No |
| 8 | Left breast | invasive ductal carcinoma | III | Neg | Neg | Neg | Yes | Yes | Yes |  |
| 9 | Left breast | Fibroadenoma | III | Neg | Neg | Neg | Yes | Yes | Yes |  |
| 10 | Left breast | invasive ductal carcinoma | II | Neg | Neg | Neg | No | Yes | No | No |
| 11 | Right breast | - | III | Neg | - | Neg | Yes | Yes | Yes | Yes |
| 12 | Left breast | invasive ductal carcinoma | III | Neg | Neg | Neg | Yes | Yes | No | No |
| 13 | Right breast | invasive ductal carcinoma | III | Neg | Neg | Neg | Yes | Yes | No | No |
| 14 | Left breast | Infiltrating ductal | III | Neg | Neg | Neg | No | Yes | Yes | No |
| 15 | Left breast | invasive ductal carcinoma | I | - | - | - | - | Yes | Yes | Yes |
| 16 | Left breast | invasive ductal carcinoma | I | - | - | - | - | No | No | No |
| 17 | Left breast | Inflammatory mammaty | III | Neg | Neg | Neg | - | No | No | Yes |
| 18 | Left breast | Infiltrating ductal | III | Neg | Neg | Neg | Yes | No | No | No |
| 19 | Right breast | Infiltrating ductal | III | Neg | Neg | Neg | Yes | No | No | No |
| 20 | Right breast | invasive ductal carcinoma | III | Neg | Neg | Neg | No | Yes | Yes | Yes |
| 21 | Left breast | Infiltrating ductal | III | Neg | Neg | Neg | Yes | Yes | Yes | No |
| 22 | Left breast | Infiltrating ductal | I | Neg | Neg | Neg | Yes | Yes | Yes | No |
| 23 | Right breast | invasive ductal carcinoma | II | Neg | Neg | Neg | No | Yes | Yes | No |
| 24 | Right breast | invasive ductal carcinoma | IV | - | - | - | Yes | Yes | Yes | Yes |
| 25 | Right breast | invasive ductal carcinoma | III | Neg | Neg | Neg | Yes | Yes | No | No |
| 26 | Right breast | invasive ductal carcinoma | III | Neg | Neg | Neg | Yes | Yes | Yes | No |
| 27 | Right breast | Infiltrating ductal | III | Neg | Neg | Neg | Yes | Yes | No | No |
| 28 | Left breast | invasive ductal carcinoma | II | Neg | Neg | Neg | Yes | Yes | Yes | Yes |
| 29 | Left breast | invasive ductal carcinoma | III | Neg | Neg | Neg | Yes | Yes | - | Yes |
| 30 | Right breast | Infiltrating ductal | II | Neg | Neg | Neg | No | Yes | - | - |
| 31 | Left breast | invasive ductal carcinoma | II | Neg | Neg | Neg | No | No | - | - |
| 32 | Left breast | invasive ductal carcinoma | I | Neg | Neg | Neg | No | No | - | - |
| ER: Estrogen receptor; PR: Progesterone receptor; Her2/neu: human epidermal growth factor receptor 2. |  |  |  |  |  |  |  |  |  |  |

### Cell Lines and cell Culture Conditions

Human Embryonic Kidney 293T (HEK293T) cell line and BC cell lines including MCF-7 and MDA-MB-231 and also an immortalized normal breast epithelial cell line, MCF-10A as control, were purchased from the International Centre for Genetic Engineering and Biotechnology (ICGEB) (Tehran, Iran). MCF-7, MDA-MB-231 and HEK293T cell lines were seeded in RPMI-1640 medium supplemented with 10 % fetal bovine serum (FBS). MCF-10A cell line was grown in DMEM:Ham’s F12 (1:1) supplemented with 5% Horse serum, 2mmol L-Glutamine, 10µg/ml Insulin, 20ng/ml Epithelial growth factor (EGF), 0.5 µg/ml Hydrocortisone and 100ng/ml cholera toxin. All cells were incubated at 37 °C in a humidified 5% CO_2_ atmosphere.

### *In-Silico* prediction of miRs

HDAC8 gene sequence was obtained from NCBI database (NG_015851.1). To predict the miRs that target the 3′UTR of HDAC8 with the highest possibility, we used the following databases: Diana-microT, RNA22, TargetScan, PicTar, miRanda, miRDB, and mirPath. At the next step, we applied mirZ database to identify miR targets with comparing their expression in different cell lines and miRCancer, Qiagen and dbDEMC 2.0 databases to provide a list of miRs in different human cancers. Finally, miRnalyze and DIANA-miRPath v3.0 databases were used to determine the role of the predicted miRs in HDAC8 signaling pathways (Table 2).

**Table 2:** Databases and algorithms for determination of target mRNAs and corresponding miRs.

|  |  |  |
| --- | --- | --- |
| Database | Algorithm | Website |
| miRanda | Complementarity | http://www.kcc.org |
| TargetScan | Seed complementarity | http://www.targetscan.org |
| TargetScanS | Seed complementarity | http://www.targetscan.org |
| DIANA microT | Thermodynamics | http://diana.pcbi.upenn.edu |
| PicTar | Thermodynamics | http://pictar.bio.nyu.edu |
| RNAHybrid | Thermodynamics | http://bibiserv.techfak.unibielefeld.de/rnahybrid |
| miTarget | SVM | http://cbit.snu.ac.kr/~miTarget |
| miTarget | validated targets | http://diana.pcbi.upenn.edu/tarbase.html |
| miRwalk | Integrative | http://www.ma.uni-heidelberg |

### RNA purification and quantitative real-time PCR (qRT-PCR) analysis

Total RNA containing small RNAs was extracted from cancerous, normal tissues and cell lines using RNX™-Plus reagent (Cinnagen, Tehran, Iran) according to the manufacturer’s instruction. cDNA was synthesized from total RNA using RevertAid First Strand cDNA Synthesis Kit (Fermentas – Thermo Scientific) according to the manual. Quantitative real-time PCR was performed with a SYBR Premix ExTaqII kit (Takara, Japan) and the Rotor-Gene 6000 apparatus (Corbett Research, Mortlake, NSW, Australia). Quantification of miRs expression was carried out using an YBR^R^ Premix ExTaq™ kit (Takara, Japan) according to the manufacturer’s instruction. *HPRT* and SNORD47 (U47) were used for mRNA and miR data normalization respectively. Primer sequences are listed in the Table 3.

**Table 3:**
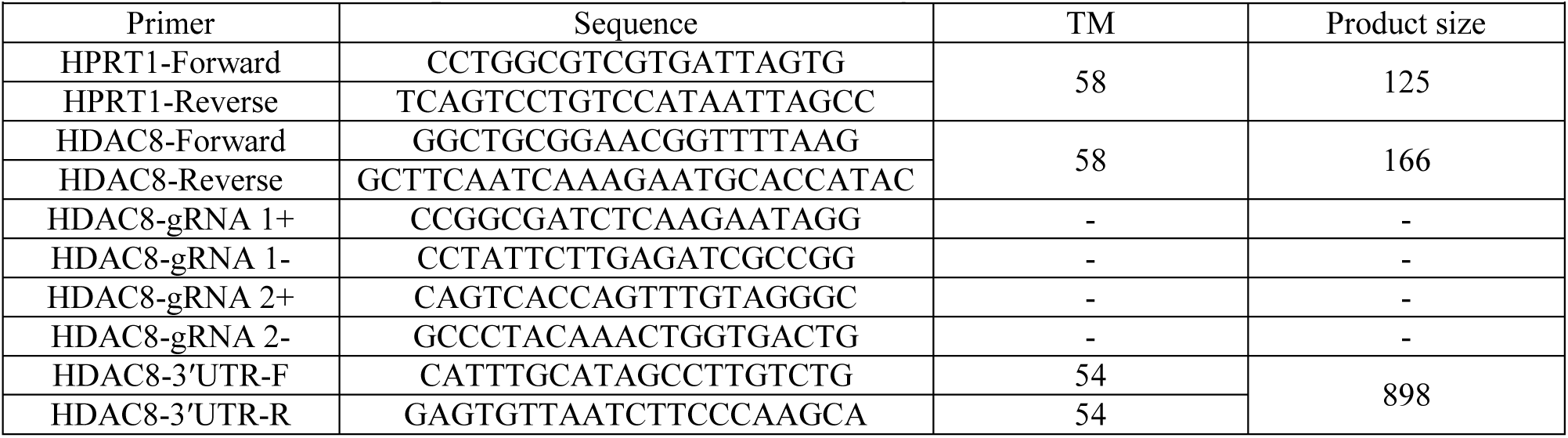
Primers used for qPCR and construction of using vectors.

### CRISPR/Cas9 mediated knockout of the *HDAC8* gene in MCF-7 and MDA-MB-231cell line

To knockout (KO) the expression of *HDAC8* in the MCF-7 and MDA-MB-231cell lines, CRISPR/Cas9 based KO strategy was applied. CRISPR sgRNAs were designed using crispr.mit.edu and the oligos were made by Macrogen Inc. (Seoul, Korea). pCAG-eCas9-GFP-U6-gRNA vector (Addgene 79145) without ITR element (Hojland Knudsen et al., 2018) was digested with BbsI enzyme for gRNA cloning. The gRNAs sequences used for knocking out of the *HDAC8* gene are illustrated in table 3. The prepared eCas9/gRNA expression vectors co-transfected to the cell lines using Lipofectamine 2000 (Invitrogen, Carlsbad, CA, USA). The eCas9 expression vector without any cloned gRNA was used for the control.

### Transient transfection

Mcf-7 and MDA-MB-231 cell lines were seeded in six-well plates at a density of 1.5 × 10^5^ cells/well and grown to 90% confluency. Then, the cells transiently transfected with 100 nM miR-216b-5p mimics, negative control and inhibitor (Exiqon). Transfection was performed by Lipofectamine 2000 according to the manufacturer’s instructions. After transfection, cells were kept in a culture medium containing 10% FBS for up to 72 h. Subsequently, total RNA and protein were extracted and used for further investigations.

### Luciferase reporter assay

The PCR purified HDAC8-3′UTR (WT-UTR) containing the predicted miR-216b-5p binding sites were cloned into the psiCHECK™-2 Vector (Promega, Madison, USA) using specific primers for HDAC8-3′UTR (Table 3). The vector was digested by SgfI and NotI restriction enzymes. A mutant luciferase vector with miR-216b-5p changed pairing site (Mut-UTR) was also constructed. Wild and mutant types of HDAC8-UTR plasmid were confirmed by DNA sequencing. When HEK293T cells reached 80% confluence in 24-well plate, cells were co-transfected with luciferase vector and miR-216b-5p mimic, inhibitor and negative control using Lipofectamine 2000. After 48 h, luciferase activities were detected by a dual-luciferase reporter assay system (Promega, Madison, US) according to the manufacturer’s instructions.

### Cell cycle analysis

Flow cytometry was applied for cell cycle assessment in different groups. Briefly, 72h after transfection, cells were collected and fixed with 70% ethanol at 1 h at 4 °C. The fixed cells washed with PBS, then incubated with propidium iodide (PI) (Sigma-Aldrich) at room temperature for 1h and samples were analyzed using a BD FACScalibur flow cytometer (BD Biosciences, Mountain View, CA).

### Cell proliferation assay

Mcf-7 and MDA-MB-231 cell lines were seeded in a 96-well plate at 5000 cells per well then transfected with miR-216b-5p mimic, negative control and inhibitor and incubated for 96 h. Cell viability was assessed after transfection using a commercial 3–2, 5-diphenyl tetrazolium bromide (MTT) assay kit (Sigma, St Louis, MO) according to the manufacturer’s instructions at designated times (6 consecutive days). The absorbance of each well was measured with a microplate reader set at 450 nM.

### Colony formation assay

Firstly, the Mcf-7 and MDA-MB-231 cell lines were transfected with miR-216b-5p mimic, negative control, inhibitor and HDAC8 knockout vector in 6-well culture dishes. The plates were covered with a layer of 0.6% agar in a medium supplemented with 20% FBS. A total of 1,000 cells were prepared in 0.3% agar and incubated for 14 days at 37 °C and 5% CO2 conditions. The resulting colonies stained with 0.04% crystal violet for 40 minutes, then rinsed with PBS again. Finally, the numbers of colonies per well were counted.

### Western Blotting

All treated and control groups were harvested using a RIPA lysis buffer supplemented with a protease inhibitor cocktail and phosphatase inhibitor cocktail (Sigma-Aldrich, USA) at 4° for 10 minutes. Protein concentrations of the cell lysates in different groups was further determined with BCA protein assay kit (Bio basic, Canada). The aliquots of 40 µg of lysates were electrophoresed by 12% SDS-PAGE and subsequently transferred onto a PVDF membrane (Millipore, USA). Non-fat milk powder (5%) was used for blocking the membranes at room temperature for 1h, then the membrane were incubated overnight with primary antibodies (Minneapolis, MN 55413). They were then incubated with horseradish peroxidase-labeled secondary antibodies (Santa Cruz, 10513) at room temperature. Finally, the signals were detected using an ECL detection kit (Sigma-Aldrich), and the membranes were scanned and analyzed using a Flowjo imaging system with imaging software (version quantity 1). Protein expression was normalized to an endogenous reference (GAPDH, Santa Cruz, D1717) and relative to the control. A protein ladder (Fermentas, Republic of Lithuania) was used as a molecular marker.

### Statistical analysis

Statistical analyses were performed using SPSS 16 (SPSS Inc., Chicago, USA). Results were presented as Mean ± SD. Comparison of the possible differences between studied groups were done by the independent samples T test. One Way ANOVA followed by Post Hoc, Tukey, and Dunnett tests were used to analyze mean differences between more than two groups. The association between two variables was calculated using the Spearman correlation coefficient. In all performed hypothesis tests, a P value less than 0.05 was considered as statistically significant.

## Results

### Expression of HDAC8 and miR-216b-5p in breast cancer cell lines and clinical specimens

To analyze the clinicopathologic status, we detected the expression of HDAC8 and miR-216b-5p in 32 primary breast cancer tissues and 32 normal adjacent tissues using qRT-PCR. The expression of miR-216b-5p in cancerous tissues were significantly decreased in compared with adjacent normal tissues (0.0024±0.00025 (r.u.) *vs*. 0.004±0.00044 (r.u.), respectively) (Figure 1A and B) (*p* value = 0.002). Our results also revealed that miR-216b-5p expression is decreased in metastatic breast cancer cell line MDA-MB-231 compared to non-metastatic (MCF-7) and control (MCF-10A) cell lines (0.00095±0.00001 (r.u.), 0.0048±0.00008 (r.u.) and 0.0062±0.0001 (r.u.), respectively) (Figure 1C). On the other hand, we found that HDAC8 expression level was highly increased in breast cancer tissue samples than normal adjacent tissues (4.86±0.44 (r.u.) *vs*. 3.52±0.43 (r.u.), respectively) (Figure 1D and E) (*p* value = 0.034). Besides, our results showed that the HDAC8 was also up-regulated in breast cancer (MDA-MB-231 and MCF-7) cell lines as compared to control (MCF-10A) cell line (0.32±0.0092 (r.u.), 0.29±0.01 (r.u.) and 0.1±0.0097 (r.u.), respectively) (Figure 1F). Finally, linear regression analysis confirmed that there was a negative statistically significant correlation between HDAC8 and miR-216b-5p in clinical specimens and cell lines (*r^2^* = −0.3952, *p* value = 0.037, Figure 1G).

**Figure 1:**
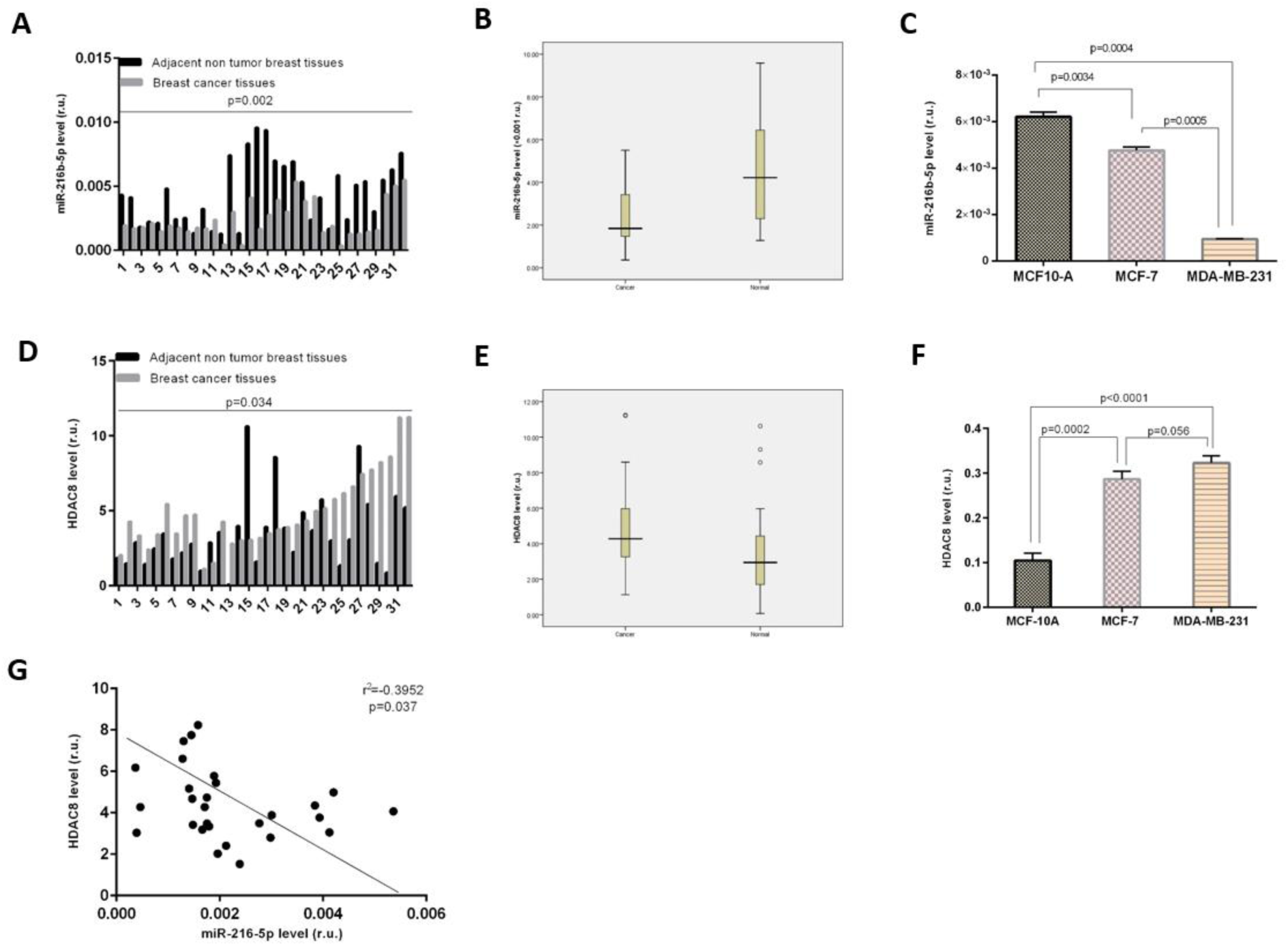
Relative expression level of miR-216b-5p and HDAC8 in breast cancer tissues and matched normal adjacent cancer tissues: 1A: miR-216b-5p level (in r.u.) separately illustrated in studied subjects. 1B: miR-216b-5p level (in r.u.) are indicated as a gray value was in a lower concentration (2.4±0.25 (×10^-3^r.u.) for the cancer group and increasing concentration to 4±0.44 (×10^-3^r.u.) for normal tissues with a significant *p* value=0.002 for their difference. 1C: miR-216b-5p levels (in relative units) are indicated as a column value of 100% for the MCF-10A control cell line and decreasing percentage to 77% (0.23 fold decrease) for non-metastatic MCF-7 cell line and lower to 15.32% (0.85 fold decrease) for the metastatic MBA-MD-231 cell line with a lower significant *p* value of 0.0004 for their difference. 1D: HDAC8 level (in r.u.) separately illustrated in studied subjects. 1E: HDAC8 level (in r.u.) are indicated as a gray value was in a higher concentration (4.86±0.44 (r.u.)) for the cancer group and decreasing concentration to 3.52±0.43 (r.u.) for normal tissues with a significant *p* value=0.034 for their difference. 1F: HDAC8 levels (in relative units) are indicated as a column value of 100% for the MCF-10A control cell line and increasing percentage to 290% (0.29 fold increase) for non-metastatic MCF-7 cell line and higher to 320% (0.32 fold increase) for the metastatic MBA-MD-231 cell line with a lower significant *p* value <0.0001 for their difference. 1G: Linear regression analysis confirmed that tissue level of HDAC8 expression inversely increased (r^2^=-0.3952, *p* value=0.037) by decreasing of miR-216b-5p expression.

### Diagnostic value of HDAC8 and miR-216b-5p as potential tumor marker in breast cancer

The means and standard deviations of HDAC8 for cancerous and normal tissues were 4.86±0.44 (r.u.) and 3.52±0.43 (r.u.), respectively. The corresponding values for miR-216b-5p were 0.0024±0.00025 and 0.004±0.00044 in BC and healthy specimens, respectively. Using a cut-off level of 3.47 (r.u.) for HDAC8, sensitivity and specificity were 69% and 60% respectively. The corresponding values for miR-216b-5p, using a cut-off level of 0.0020 (r.u.), were 81% and 60%. Areas under the curve (AUC) for HDAC8 and miR-216b-5p were 0.687 and 0.752 respectively. The Positive Predictive Value (PPV) and Negative Predictive Value (NPV), likelihood ratio (positive and negative) for both HDAC8 and miR-216b-5p are shown in Table 4. Besides, our results showed that decreased level of miR-216b-5p was significantly associated with tumor size and lymph node invasion (Spearman’s *ρ*= −0.382, *p* value=0.041 and Spearman’s *ρ*= −0.373, *p* value=0.039). There was not any significant association between HDAC8 tissue level and clinical outcomes (Table 5).

**Table 4:** Diagnostic values for HDAC8 and miR-216b-5p in breast cancer.

| **Table 4: Diagnostic values for HDAC8 and miR-216b-5p in breast cancer** |  |  |  |  |  |  |  |  |
| --- | --- | --- | --- | --- | --- | --- | --- | --- |
|  | Cut-off point (r.u.) | Sensitivity | Specificity | PPV | NPV | LR+ | LR- | AUC |
| HDAC8 | 3.47 | 69% | 60% | 0.61 | 0.64 | 1.73 | 0.52 | 0.687 |
| miR-216b-5P | 0.0020 | 81% | 60% | 0.67 | 0.76 | 2.03 | 0.32 | 0.752 |
| LR = Likelihood ratio; PPV = Positive predictive value; NPV = Negative predictive value; AUC= Area under the curve; HDAC8= Histone deacetylase 8; miR-216b-5p= microRNA-216b-5p. |  |  |  |  |  |  |  |  |

**Table 5:** Correlation of miR-216b-5p and HDAC8 with clinical characteristics of the studied subjects.

|  |  |  |  |  |  |  |
| --- | --- | --- | --- | --- | --- | --- |
|  |  | Tumor size | Grade | Lymph node invasion | Vascular invasion | Necrosis |
| miR-216b-5p | Spearman's $\rho$ correlation coefficient | -0.382 | -0.287 | -0.373 | 0.057 | -0.289 |
| | $p$ value | 0.041 | 0.112 | 0.039 | 0.777 | 0.136 |
| HDAC8 | Spearman's $\rho$ correlation coefficient | -.069 | -0.051 | -0.143 | -0.067 | -0.099 |
| | $p$ value | 0.722 | 0.783 | 0.443 | 0.741 | 0.615 |
| HDAC8= Histone deacetylase 8; miR-216b-5p= microRNA-216b-5p. |  |  |  |  |  |  |

### Effects of ectopic expression of miR-216b-5p on cell proliferation and cell cycle

To study the role of miR-216b-5p in BC cells, MCF-7 and MDA-MB-231 cell lines were transiently transfected with miR-216b-5p mimics and inhibitor, then the following overexpression or inhibition of miR-216b-5p by mimics or inhibitor were detected by qRT-PCR. As shown in figures 2A and B, after transfection of BC cell lines with miR-216b-5p mimics and inhibitor, expression of miR-216b-5p significantly increased (0.71±0.015 and 0.14±0.012 (r.u.) for MCF-7 and MDA-MB-231, respectively) (*p* value<0.0001) and decreased (0.015±0.001 and 0.0006±0.00008 (r.u.) for MCF-7 and MDA-MB-231, respectively) (*p* value <0.0001 and =0.01 for MCF-7 and MDA-MB-231, respectively), compared with NC and mock controls (0.035±0.002 and 0.001±0.0001 (r.u.), 0.041±0.004 and 0.0009±0.00008 (r.u.) for MCF-7 and MDA-MB-231, respectively). We used the MTT assay for evaluating the proliferation status of BC cell lines after transfection with miR-216b-5p mimics and inhibitors. As shown in figures 2C and D the miR-216b-5p mimics significantly inhibited proliferation of MCF-7 and MDA-MB-231 cell lines by 11% and 21% (*p* value= 0.008 and 0.006, respectively), 16% and 36% (*p* value= 0.004 and 0.0005, respectively) and 27% and 43% (*p* value= 0.002 and 0.0001, respectively) at 48, 96 and 144 h, respectively, as compared to the controls. On the other hand, miR-216b-5p inhibitor transfection in MCF-7 and MDA-MB-231 cell lines enhanced cell proliferation (Figures 5A and B). We next studied the growth of the cells using a flow cytometric assay of the cell distribution. Figures 6A and B show the function of miR-216b-5p on cell cycle profile of MCF-7 and MDA-MB-231 cell lines at 72 h. Accordingly, overexpression of miR-216b-5p resulted in G0/G1 phase arrest in MCF-7 and MDA-MB-231 cell lines after 72 h of exposure compared with NC and mock controls, while cells treated with miR-216b-5p inhibitors showed a significant decrease in G0/G1 arrest as compared to controls (Figures 6A and B). Finally, we further inspected the effects of miR-216b-5p on BC cells clonogenicity by a colony formation assay. Our results clearly showed that the miR-216b-5p-transfected MDA-MB-231 and MCF7 cells formed smaller amounts of colonies per well. A significant reduction in colony formation was detected in MCF-7 and MDA-MB-231 cells transfected with miR-216b-5p mimic (Figures 7A and B) while miR-216b-5p inhibitor transfection suppressed this effect (*p* Value < 0.0001).

**Figure 2:**
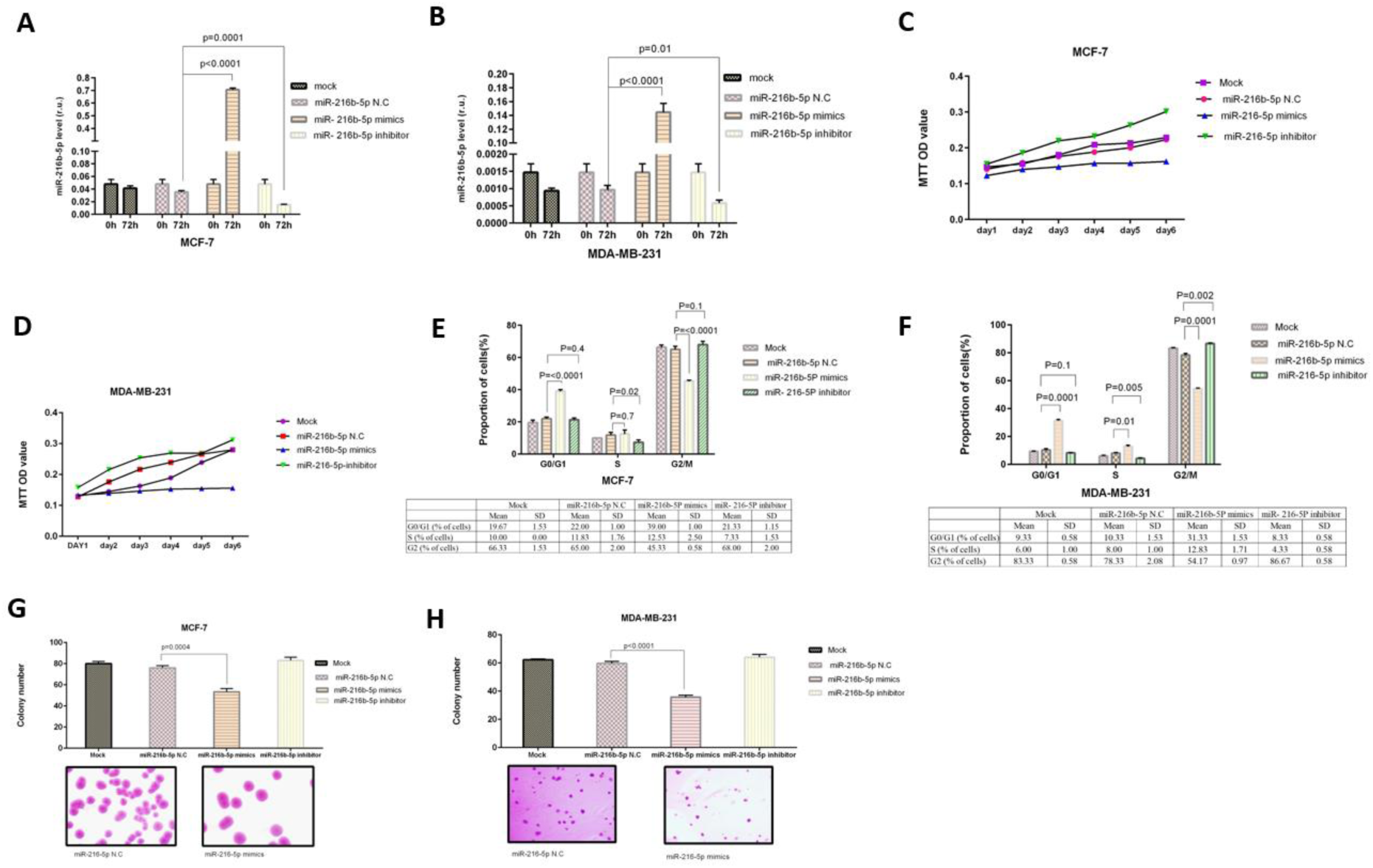
Overexpression of miR-216b-5p inhibited cell proliferation and colony formation and induced G0/G1 arrest in BC cell lines. 2A and B: The relative level of miR-216b-5p after transfection with mimics inhibitor in MCF-7 (2A) and MDA-MB-231(2B) cell lines. 2C and D: MTT assay for 6 days of and MDA-MB-231 (2D) cells transfected with miR-216b-5p mimics (*p* value=0.002, 6D) for MCF-7 (2C) and (*p* value=0.0001, 6D) for MDA-MB-231 as compared to NC. 2E and F: BC cell lines transfection with miR-216b-5p inhibited the entry of MCF-7 (2E) and MDA-MB-231 (2F) cells into G2/M phase. 2G and H: Representative images of the colony formation assay show that, following treatment with miR-216b-5p, MCF-7 (2G) and MDA-MB-231 (2H) cells formed fewer colonies compared with controls. The number of clones formed by the MCF-7 (2G) and MDA-MB-231 (2H) cell lines were derived from the colony formation assay. Results are presented as the mean ± standard deviation from three independent experiments. *p* value=0.0004 and <0.0001 for MCF-7 and MDA-MB-231 compared with the NC.

### HDAC8 is involved in miR-216b-5p -regulated proliferation, cell cycle arrest and colony formation inhibition in breast cancer cells

To identify whether HDAC8 involves in miR-216b-5p-suppressed breast cancer progression, HDAC8 was knocked out by CRISPR/Cas9 method in MDA-MB-231 and MCF7 cell lines and BC cellular functions including proliferation, cell cycle and colony formation were checked on the knocked out selected population of the cell lines. Decline in HDAC8 expression after transfection with knockout vector were detected in RNA and protein levels using qRT-PCR and Western blot methods, respectively. qRT-PCR analysis of HDAC8 expression showed significant decrease in HDAC8 expression levels after transfection of BC cell lines with HDAC8 knockout vector compared to empty vector and mock controls (0.096±0.005 and 0.14±0.004 (r.u.), respectively) (Figures 3A and C). Besides, the corresponding protein levels also confirmed the highly decline in HDAC8 in knockout group as compared to controls (0.76±0.03 and 0.63±0.04, respectively) (Figures 3B and D). As shown in figure 3C and D, in MDA-MB-231 loss of function of the HDAC8 in gene and protein levels had higher significance value than MCF-7 cell lines. After transfection of MDA-MB-231 cell lines with HDAC8 knock out vector, the expression of HDAC8 in gene and protein level were extremely down-regulated (0.019±0.0005 (r.u.), 0.8±0.14, respectively) when compared to empty vector group (0.14±0.006 (r.u.), 1.1±0.0005, respectively) and mock group (0.17±0.006 (r.u.), 1.1±0.1, respectively) (*p* value<0.0001, *p* value=0.04). Next the capacity of cell growth was confirmed by flowcytometry and cologenicity tests. Cell cycle analysis revealed that the percentage of cells at G1/G0 phase dramatically increased from 21% to 50% and 10% to 24% in MCF-7 and MDA-MB-231 cell lines after transfection with HDAC8 KO vector at 72 h after transfection (Figures 3E and F) compared to empty vector. In addition, we used colony formation assay to investigate the role of HDAC8 on clonogenic survival, and results demonstrated that knocking out of HDAC8 caused a decrease in the clonogenic survival of MCF-7 and MDA-MB-231 cells (46±5 and 26±2.5 colonies, respectively) compared with empty vector (73±3 and 51±2 colonies, respectively) and mock group (78±9 and 56±2.1 colonies,respectively) (*p* value= 0.0035 and 0.0003, respectively). (Figures 3G and H)

**Figure 3:**
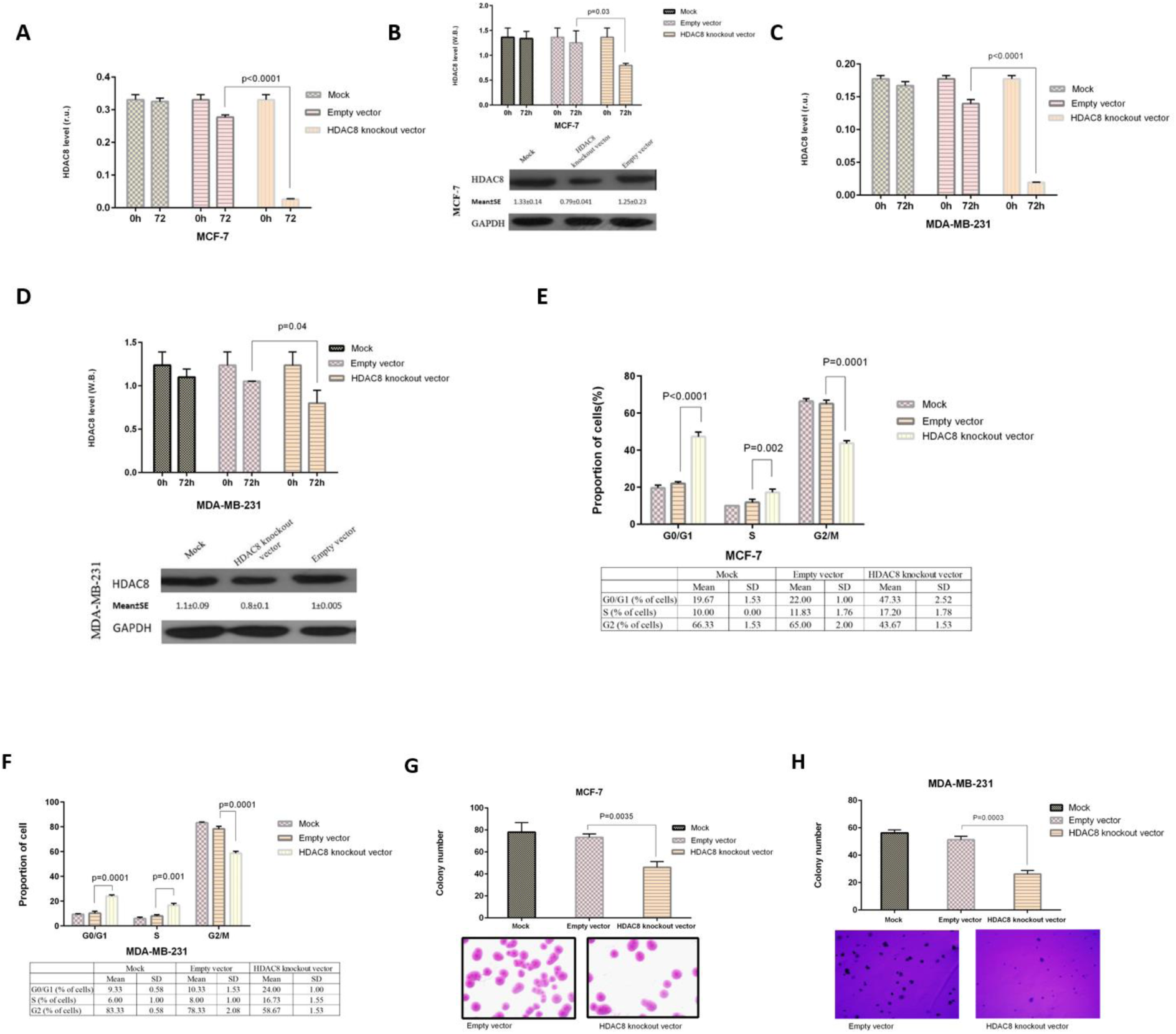
Down-regulation of HDAC8 inhibited cell proliferation and colony formation and induced G0/G1 arrest in BC cell lines. 3A and B: The relative expression of HDAC8 extremly decreased in gene (3A) and protein (3B) level after transfection with HDAC8 knockout vector in MCF-7 cell line. 3C and D: Transfection of MDA-MB-231 cell line with HDAC8 knockout vector also resulted in highly decline in HDAC8 expression in gene (3C) and protein (3D) level. 3E and F: BC cell lines transfection with HDAC8 knockout vector inhibited the entry of MCF-7 (3E) and MDA-MB-231 (3F) cells into G2/M phase. 3G and H: Representative images of the colony formation assay show that, following treatment with HDAC8 knockout vector, MCF-7 (3G) and MDA-MB-231 (2H) cells formed fewer colonies compared with controls. The number of clones formed by the MCF-7 (3G) and MDA-MB-231 (3H) cell lines were derived from the colony formation assay. Results are presented as the mean ± standard deviation from three independent experiments. *p* value=0.0035 and 0.0003 for MCF-7 and MDA-MB-231 compared with the NC.

### miR-216b-5p directly targets 3′UTR of HDAC8 mRNA

In order to search for the effective functional target of miR-216b-5p and its mechanism we used online computational miRNA target prediction algorithm (TargetScan 6.0). This algorithm predicted the 3′-UTRs of HDAC8 mRNA contain putative miR-216b-5p binding sites. We evaluated three different predicted miR-216b-5P binding sites that were found in 3′UTR region of HDAC8 gene (Figure 4A). We then investigated the mechanism by which miR-216b-5p suppresses the progression of breast cancer tumors. To confirm whether HDAC8 is a direct target of miR-216b-5p, we inserted the wild-type and the 550bp-mutant form of HDAC8 3′-UTR into the psiCHECK-2 vector at the downstream of the Renilla luciferase coding sequence separately. As illustrated in figure 4B miR-216b-5p rather than control significantly suppressed the Renilla luciferase activity in HEK293T cells, which was compromised when the binding site of miR-216b-5p was mutated. Furthermore, we found that overexpression of miR-216b-5p reduced the endogenous HDAC8 protein expression in both MCF-7 and MDA-MB-231 cells, whereas transfection with miR-216b-5p inhibitor increased the level of HDAC8 mRNA in MCF-7 and MDA-MB-231 cells. The results were confirmed by Western blot analysis (Figures 4C-F).

**Figure 4:**
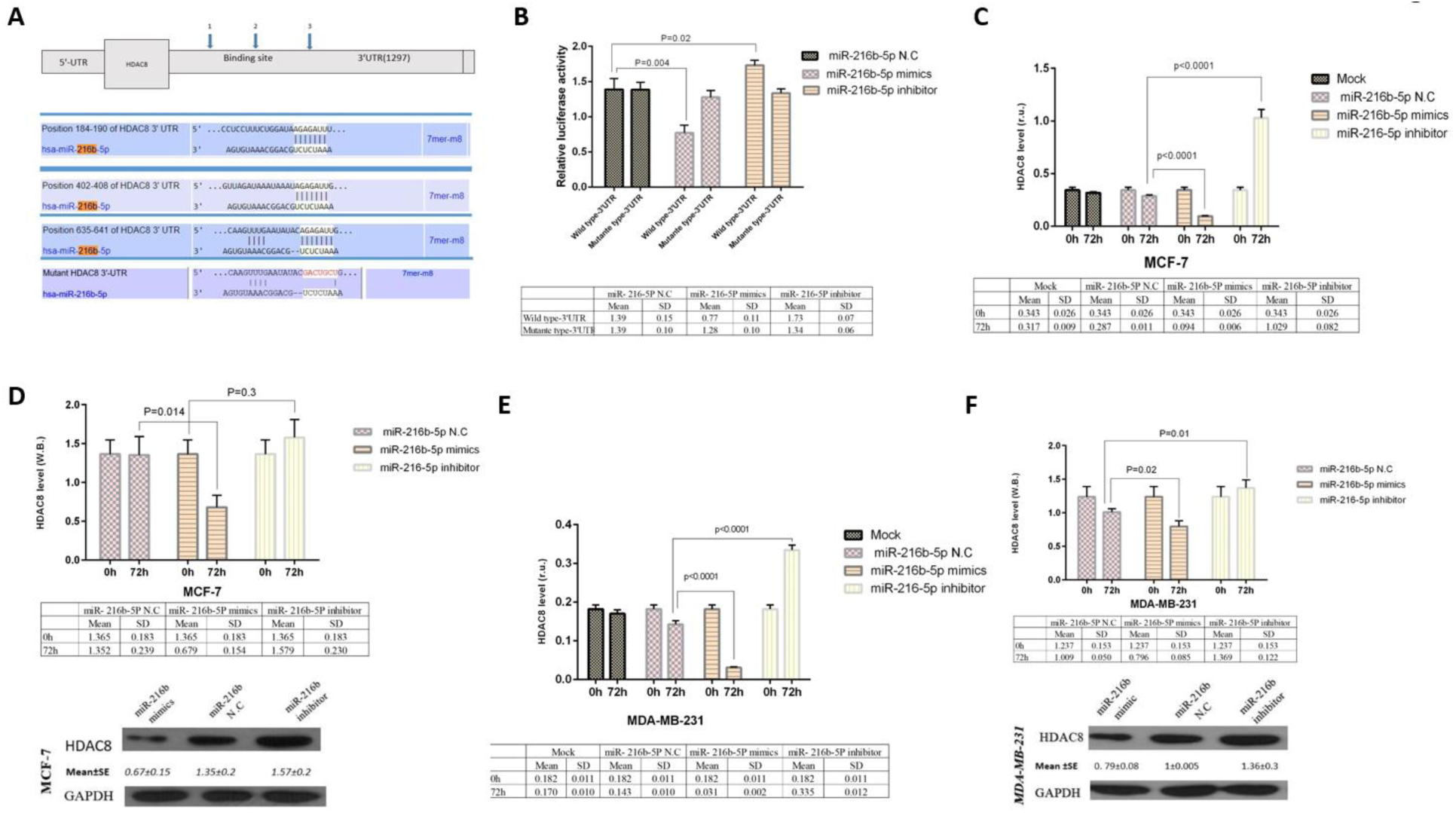
miR-216b-5p directly targets the 3′-UTR of HDAC8 gene. 4A: *In-silico* predicted of the pairing of miR-216b-5p to sites in the 3′-UTR of the human HDAC8 gene. 4B: Relative luciferase activity of the HDAC8 wild type-3′UTR and mutant type-3′UTR luciferase constructs in HEK293T cell line transfected with miR-216b-5p mimics, inhibitor or NC. 4C and D: The relative expression of HDAC8 extremly decreased in gene (4C) and protein (4D) level after transfection with miR-216b-5p mimics in MCF-7 cell line. 4E and F: Transfection of MDA-MB-231 cell line with miR-216b-5p mimics also resulted in highly decline in HDAC8 expression in gene (4E) and protein (4F) level.

## Discussion

In the present study, we examined the functional significance of HDAC8 and miR-216b-5p in breast cancer. In this study we showed that miR-216b-5p is significantly down-regulated in human breast cancer tissue and breast cancer cell lines and has a negative correlation with the HDAC8 expression level. Bioinformatics analyses identified various tumor suppressive miRs that can target the 3′-UTR of HDAC8 and among them we showed that the expression of miR-216b-5p is highly correlated with the expression of HDAC8 in different BC cell lines. Hence, we report miR-216b-5p to be one of the most potent miR targeting HDAC8. Our results demonstrate that miR-216b-5p down regulates the expression of HDAC8 by directly binding to its 3′-UTR sequence. In addition, a significant decrease in the HDAC8 expression level was observed upon miR-216b-5p overexpression. Also, a significant down-regulation of HDAC8 expression was seen while the miR-216b-5p mimics transfected to the cells.

Although the prognosis of breast cancer has improved due to advances in treatment concepts and treatment methods, the benefit of current approaches is limited given the emergence of resistance to treatments (Jiang et al., 2014). The beginning and development of cancer, conventionally seen as a genetic disease, has been determined to involve epigenetic abnormalities in addition to genetic alterations (Shen et al., 2016). The epigenetic mechanisms of cancer consist of DNA methylation, histone modification, nucleosome positioning, and noncoding RNA expression, specifically microRNA expression (Shen et al., 2016). Recent studies revealed the direct correlation between alterations in histone proteins and breast tumorigenesis (Connolly and Stearns, 2012; Kanwal et al., 2015; Koeneke et al., 2015). In breast cancer, HDAC8 is highly over expressed in triple-negative breast cancer (Hsieh et al., 2016) and responsible for late stage particularly in triple-negative breast cancer, poor prognosis and poor treatment response (Hsieh et al., 2016). Therefore regulation of HDAC8 at the protein level can play a critical role in tumor development. Recently, it has been more attention to targeted inhibition of HDAC8 as a potential cancer cells growth suppressor in vitro and in vivo (Rettig et al., 2015).

In line with previous studies, we showed that HDAC8 is overexpressed in clinical specimens and breast cancer cell lines. Furthermore, to confirm the oncogenic activity of HDAC8 in breast cancer we performed the knocking out of the HDAC8 using the CRISPR/Cas9 method and found out that by partial knocking out of the endogenous HDAC8 at the cells population level inhibit the oncogenic functions of HDAC8.

Altered expressions of different miRNAs have been also reported in breast cancer pathogenesis (Bischoff et al., 2014; Liang et al., 2014). For better therapeutic approach and better understanding of the molecular mechanism of cancer growth, the roles of miRs in cancer progression have been studied extensively.

In human cancer growth and metastasis miR-216b-5p takes a big role as a tumor suppressor in nasopharyngeal carcinoma (Deng et al., 2011), colorectal cancer (Kim et al., 2012), breast cancer (Zheng et al., 2014) and hepatocellular carcinoma (Liu et al., 2015). On the other hand it also regulates the apoptosis by mediating the activity of c-Jun (Xu et al., 2016) and Autophagy (Chen et al., 2016). However the role of miR-216b-5p for the regulation of HDAC8 has not been studied yet. Our result, for the first time, explain that the miR-216b-5p function as a tumor suppressor and inhibit the proliferation, cell growth and colony formation by regulating the expression of HDAC8 in breast cancer.

Despite diagnostic and therapeutic advances during the last decades, unsuccessful chemotherapy because of fail to respond or resistance to chemotherapeutic agents are still remain in about 50% of the BC patients. This has caused to BC still remains as the second death-leading cause from cancer among women. Drug resistance is still a major clinical problem to successful treatment in breast cancer patients. In addition to the chemoresistance, the metastasis of cancerous cells in patients with solid tumors likes those created from breast tissue, is responsible for about 90% of deaths. As a result there is more attention to development of a new therapeutic target for the treatment of BC patients. By these observations we found that miR-216b-5p may be a potent therapeutic molecule for the treatment of metastatic breast cancer by regulating the activity of HDAC8.

In conclusion, our study gives the idea of molecular mechanism involved in metastatic breast cancer growth via dysregulation of miR-216b-5p resulted HDAC8 overexpression. Down-regulation of HDAC8 expression in breast cancer suppresses cell proliferation, cell cycle arrest and colony formation. HDAC8 suppression via the miR-216b-5p possibly open a new therapeutic approach to treat the metastatic breast cancer.

## Acknowledgments

The author wish to thank all patients and health stuffs who participated in this study. Financial support from Kurdistan University of medical sciences is highly appreciated.

## Disclosure of potential conflicts of interest

Mr MN Menbari declares no potential conflicts of interest with respect to the research, authorship, and/or publication of this article. Dr K Rahimi declares that he has no conflict of interest. Dr A Ahmadi declares that he has no conflict of interest. Dr S Mohammadi-Yeganeh declares that she has no conflict of interest. Dr A Elyasi declares that he has no conflict of interest. Mrs. N Darvishi declares that she has no conflict of interest. Dr V hosseini declares that she has no conflict of interest. Dr M Abdi has received research grants from Kurdistan University of medical sciences.

## Ethical approval

All procedures performed in studies involving human participants were in accordance with the ethical standards of the ethics committee of Kurdistan University of Medical Sciences and with the 1964 Helsinki declaration and its later amendments or comparable ethical standards.

## Informed consent

Informed consent was obtained from all individual participants included in the study.

## Funding source

This work was supported by a research grant from Kurdistan University of medical sciences (Grant/Award Number: ‘IR.MUK.REC.1395/279’).

## Financial disclosure

The author has no financial relationships relevant to this article to disclose.

## Author Contributions

All authors contributed equally in this work.

